# Using Optimal F-Measure and Random Resampling in Gene Ontology Enrichment Calculations

**DOI:** 10.1101/218248

**Authors:** Weihao Ge, Zeeshan Fazal, Eric Jakobsson

## Abstract

**Background:** A central question in bioinformatics is how to minimize arbitrariness and bias in analysis of patterns of enrichment in data. A prime example of such a question is enrichment of gene ontology (GO) classes in lists of genes. Our paper deals with two issues within this larger question. One is how to calculate the false discovery rate (FDR) within a set of apparently enriched ontologies, and the second how to set that FDR within the context of assessing significance for addressing biological questions, to answer these questions we compare a random resampling method with a commonly used method for assessing FDR, the Benjamini-Hochberg (BH) method. We further develop a heuristic method for evaluating Type II (false negative) errors to enable utilization of F-Measure binary classification theory for distinguishing “significant” from “non-significant” degrees of enrichment.

**Results:** The results show the preferability and feasibility of random resampling assessment of FDR over the analytical methods with which we compare it. They also show that the reasonableness of any arbitrary threshold depends strongly on the structure of the dataset being tested, suggesting that the less arbitrary method of F-measure optimization to determine significance threshold is preferable.

**Conclusion:** Therefore, we suggest using F-measure optimization instead of placing an arbitrary threshold to evaluate the significance of Gene Ontology Enrichment results, and using resampling to replace analytical methods

## Background

Gene Ontology (GO) enrichment analysis is a powerful tool to interpret the biological implications of selected groups of genes. The gene lists from experiments such as microarrays, are gathered into clusters associated with biological attributes, and defined as GO terms^1^. The GO terms are arranged in an acyclic tree structure from more specific to more general descriptions, including biological process (BP), cellular component (CC), and molecular function (MF). GO aspires to create a formal naming system to define the biologically significant attributes of genes across all organisms. Each enriched GO term derived from a list of genes is evaluated by its significance level, i.e. the probability that the measured enrichment would be matched or exceeded by pure chance.

Enrichment tools have been developed to process large gene lists to generate significantly enriched ontologies. Huang *et.al* (2009) summarizes the tools widely used for GO enrichment^2^. Different tools emphasize different features. Gorilla^3^, DAVID^4^, g:profiler^5^ are web interfaces that integrate functional annotations including GO annotations, disease and pathway databases etc. Blast2GO^6^ extends annotation of gene list to non-model organisms by sequence similarity. GO-Miner^7^, Babelomics^8^, FatiGO^9^, GSEA^10 11^, and ErmineJ^12^ apply resampling or permutation algorithms on random sets to evaluate the number of false positives in computed gene ontologies associated with test sets. DAVID ^4^ and Babelomics ^8^ introduced level-specific enrichment analysis; that is, not including both parents and children terms. TopGO contains options, “eliminate” and “parent-child”, which eliminate or reduce the weight of genes in the enriched children terms when calculating parent term enrichment^13^. TopGO^14^ and GOstats^15^ provide R-scripted tools for ease of further implementation. Cytoscape plugin in BinGO ^16^ is associated with output tree graphs.

To calculate raw *p*-values for GO enrichment without multiple hypothesis correction, methods used include exact or asympotic (i.e. based on the hypergeometric distribution or on Pearson’s distribution), one- or two-sided tests ^17^. Rivals *et. al*. discussed the relative merits of these methods^17^. Generally, inference of the statistical significance of observed enrichment of categories in gene ontology databases can’t be assumed to be parametric, because there is no *a priori* reason to postulate normal distributions within gene ontology terms. Randomization methods are powerful tools for testing nonparametric hypotheses^18^. However, heuristic methods for testing nonparametric hypotheses have long been widely used due to lack of adequate computational resources for randomization tests. In gene ontology enrichment, a widely-used heuristic method is that of Benjamini and Hochberg^19^. In their original paper, Benjamini and Hochberg tested their method against a more computationally intensive resampling procedure for selected input data and found no significant difference, Thus the more computationally efficient Benjamini-Hochberg method was justified.

Benjamini-Hochberg has been widely applied in enrichment tools such as BinGO^16^, DAVID^4^, GOEAST^20^, Gorilla^3^, and Babelomics^8^, to name a few. The similar Benjamini-Yekutieli method is included in the GOEAST package which enables to control the FDR even with negatively correlated statistics^20 21^. A recent approach published by Bogomolov, *et.al.* (2017) deals with multiple hypothesis correction and error control for enrichment of mutually dependent categories in a tree structure using a hierarchical Benjamini-Hochberg-like correction^22^. Gossip provides another heuristic estimation of false positives that compares well with resampling in the situations tested^23^.

A randomized permutation method for assessing false positives is embedded in the protocol of Gene Set Enrichment Analysis (GSEA)^10^. Kim and Volsky^24^ compared a parametric method (PAGE) to GSEA and found that PAGE produced significantly lower *p*-values (and therefore higher putative significance) for the same hypotheses. They suggest that PAGE might be more sensitive because GSEA uses ranks of expression values rather than measured values themselves. However, they do not demonstrate that the hypothesis of normal distributions in gene ontology databases that underlies PAGE is generally true.

Noreen^25^ considered the potential of using more widely available computer power to do exact testing for the validity of hypotheses, in order to be free of any assumptions about the sampling distributions of the test statistics, for example the assumption of normality. The essence of the more exact methods is the generation of a null hypothesis by the creation and analysis of sets of randomly selected entities (null sets) that are of the same type as the test set. Then the extent to which the null hypothesis is rejected emerges from comparing the results of conducting the same analysis on the null sets and the test set. As exemplified by the over one thousand citations of this work by Noreen, these methods have been widely adopted in many areas in which complex datasets must be mined for significant patterns, as for example in financial markets.

In the present paper we utilize a straightforward random resampling method for creation of null sets and compare resultant assessments for estimating false positives with commonly used analytical methods as applied to gene ontology enrichment analysis. We also evaluate the computational cost of this method relative to analytical methods.

In applying all the cited methods and tools, it is common to apply a threshold boundary between “significant enrichment” and “insignificance”. Such assignment to one of two classes is an example of a binary classification problem. Often such classifications are made utilizing an optimum F-measure^26^. Rhee, *et.al.* (2008) have suggested application of F-measure optimization to the issue of gene ontology enrichment analysis^27^. In the present work, we present an approach to gene enrichment analysis based on F-measure optimization, which considers both precision and recall and provides a flexible reasonable threshold for data sets depending on user choice as to the relative importance of precision and recall. We also compare a resampling method to the Benjamini-Hochberg method for estimation of FDR and use with F-measure optimization.

We also consider the argument made by Powers ^26^ that the F-measure is subject to biases, and that instead of precision and recall (the constituents of the F-measure) the constructs of markedness and informedness should be considered. Whereas precision and recall are entirely based on the ability to identify positive results, informedness and markedness give equal weight to identification of negative results. We note that the Matthews Correlation Coefficient (MCC), another well-vetted measure of significance^28^, is the geometric mean of the markedness and informedness.

Our results in this paper will suggest that resampling is preferable to analytical methods to estimate FDR, since the compute costs are modest by today’s standards and that even well-accepted and widely used analytical methods may have significant error. Our results also suggest that F-measure or MCC optimization is preferable to an arbitrary threshold when classifying results as “significant” or “insignificant”. For the particular analyses in this paper, we found no significant difference in utilizing F-measure vs. MCC. in assessing significance of results in computing enrichment in gene ontology analysis.

## Methods

### Enrichment Tool

For results reported in this study (described below), the TopGO^14^ package is implemented to perform GO enrichment analysis, using the “classic” option. In this option, the hypergeometric test is applied to the input gene list to calculate an uncorrected *p*-value.

### FDR Calculation

The empirical resampling and Benjamini-Hochberg (BH) methods are used to estimate the FDR. The *p*-value adjustment using Benjamini-Hochberg is carried out by a function implemented in the R library. http://stat.ethz.ch/R-manual/R-devel/library/stats/html/p.adjust.html The resampling method is based on the definition of *p*-value as the probability that an observed level of enrichment might arise purely by chance. To evaluate this probability, we generate several null sets, which are the same size as the test set. The genes in the null sets are randomly sampled from the background/reference list. GO enrichment analysis was carried out on both test set and null set. The average number of enriched results in the null sets would be the false positives. In all the results shown in this paper, 100 null sets were used to compute the average, unless otherwise indicated. In the pipeline, available for download in Supplementary material, the number of null sets is an adjustable parameter. The ratio of false positives to predicted positives is the FDR.

### F-measure Optimization and the Matthews correlation coefficient

To evaluate F-measure and MCC, we started with evaluating true/false positive/negatives and the metrices derived from the true/false positive/negatives. The number of “predicted positive” is the number of GO terms found at a threshold. For an analytical method such as BH, the “false positive” would be (predicted positive) multiply by FDR, which is estimated by the corrected *p*-value. For resampling, the “false positive” would be the average number of GO terms found by null sets. The “true positive” is calculated by:

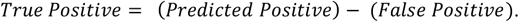

Then, we calculate the precision:

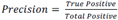

Recall is defined as

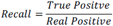

“Real Positive” is defined by

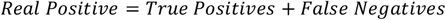

In the absence of the ability to calculate “False Negatives” directly, we estimate the number of real positives as the maximum true positive achieved across the range of possible *p*-values. This procedure is shown graphically in Figure 1 for the BH method of computing false positives, using as an example a gene list to be described in detail later in the paper. In this figure we plot predicted positives, false positives (False Discovery Rate x predicted positives), and true positives (predicted positives – false positives) vs. uncorrected *p*-value for the entire range of *p*-values from 0 to 1. At very lenient *p*-values the FDR approaches 1, resulting in the true positives approaching 0. It is difficult to evaluate false negatives and thus assign a number for “real positives”, since a false negative is an object that escaped observation, and thus can’t be counted directly. Yet such estimation is essential to applying F-measure. In our case, if we follow the trajectory of the true positives in Figure 1 as the threshold is relaxed, we see that at very stringent *p*-values all positives are true positives. As the threshold is relaxed further, more false positives are generated, so the predicted positive and true positive curves start to diverge. At *p* = 0.13 (a far higher value than would ordinarily be used as a cutoff) the true positives reach a maximum, and the number of true positives starts to decline as *p* is further relaxed. We utilize this maximum value as the maximum number of GO categories that can be possibly regarded as enriched in the data set; i.e., the number of real positives.

**Figure 1.**
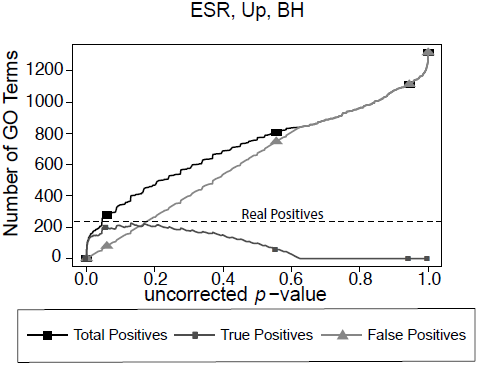
Number of positives for the yeast environmental stress response (ESR) set over the full range of uncorrected *p*-values from 0 to 1. “Predicted positives” is the number of Biological Process GO categories returned as a function of the *p*-value threshold for significance. “False Positives” is the number of predicted positives multiplied by the False Discovery Rate as calculated by the Benjamini-Hochberg formulation. “True Positives” is “Predicted Positives” minus “False Positives”. “Real Positives”, necessary to estimate number of false negatives, is estimated as the largest number of true positives computed at any uncorrected *p*-value.

Based on precision and recall at each raw *p*-value cut-off, we can obtain a table and curve of F-measure vs uncorrected *p*-value. The F_1_-measure is an equally weighted value of precision and recall. A generalized F-measure introducing the parameter β can be chosen based on the research question, whether minimization of type I (false positive) or type II (false negative) error, or balance between the two, is preferred, according to the equation:

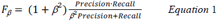

The larger the magnitude of β the more the value of *F*_*β*_ is weighted towards recall; the smaller the value of β the more the value of *F*_*β*_ is weighted towards precision. Optimizing F-measure provides us a threshold which emphasize precision (β<1) or recall (β>1), or balance of both (β=1). Note that precision and recall are extreme values of F-measure; that is, Precision=F_0_ and Recall=F_∞_.

To compare the different thresholds, we also calculated for each of them the Matthews correlation coefficient (MCC) ^28^. Originally developed to score different methods of predicting secondary structure prediction in proteins, the MCC has become widely used for assessing a wide variety of approaches to binary classification, as exemplified by the 2704 citations (at this writing) of the original paper. Perhaps even more telling, the citation rate for the seminal MCC paper has been increasing as the method is being applied in a greater variety of contexts, reaching 280 citations in 2017 alone.

In the expression below for the MCC, the True Negative (TN) is estimated using total number of GO categories in the database minus predicted positive and false negative.

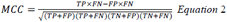

The MCC can be expressed in an equivalent expression using definition of informedness and markedness, which includes precision and recall, as well as the inversed precision and recall evaluating the proportion of true negatives:

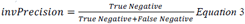

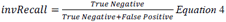

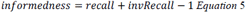

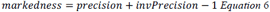

Combining equations 2-6 and some algebra we find:

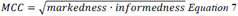

In an analogous fashion to the manner in which the F-measure may be generalized to weight either precision or recall more strongly by a variable β, so also the MCC can be generalized to more strongly weight either markedness or informedness by the expression

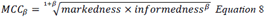

### Data Sets

#### Environmental Stress Response (ESR)

First dataset is the Yeast Environmental Stress Response (ESR) data ^29^, a robust data set for a model organism. The ESR set is list of genes commonly differentially expressed in response to environmental stresses such as heat shock, nutrient depletion, chemical stress, etc. Approximately 300 genes are up-regulated, and 600 genes are down-regulated in the ESR set. We expect this set to be “well-behaved” (give reasonable results with standard methods of analysis), since the data come from a very well annotated model organism subject to a widely studied experimental intervention.

#### Alarm Pheromone (AP)

The second data set is comprised of human orthologs to the honey bee Alarm Pheromone set^30^. The Alarm Pheromone set is a list of genes differentially expressed in honey bee brain in response to the chemical alarm pheromone, which is a component of the language by which honey bees communicate with each other. Previous studies have shown that the Alarm Pheromone set is enriched in placental mammal orthologs, compared to other metazoans including non-social insect orthologs^31^. The Alarm Pheromone set is much smaller than the ESR set, with 91 up-regulated genes and 81 down-regulated genes. We expect the AP set to be not so “well-behaved” compared to the ESR set, as we are using model organism orthologs (human) to a non-model organism (honey bee) and the organisms diverged about 600 million years ago.

#### Random Test Sets

To generate a baseline of the analysis for each data set using different FDR calculation methods, we have applied the pipeline to analyze randomly-generated sets as “test” set inputs, where FDR should equal to 1 for all uncorrected *p*-values.

The BH FDR curves are calculated in the following way: The R program p.adjust is applied to generate a list of analytically calculated FDR (BH) corresponding to uncorrected *p*-values for each “test” sets. Then the lists of FDRs are merged and sorted by uncorrected *p*-values. The FDRs are smoothed by a “sliding window” method: at each uncorrected *p*-value point, the new FDR is the average value of 11 FDRs centered by the uncorrected *p*-value point.

The Resampling FDR curves are calculated in the following way: The output uncorrected *p*-values are binned in steps of 1E-4. The counts below the upper bound of each *p*-value bin for the “test” set enrichment categories are the “Predicted positives”, and average counts for the null set enrichment categories are the “False Positives”. The process is repeated for the multiple “test” sets, and corresponding to each test set, 100 null sets were generated for “False Positive” calculation. Then the number of total and false positives are averaged, respectively. The FDR would be the quotient of the averaged total and false positives. Then, all the FDRs are plotted against the uncorrected *p*-values.

## Results

In this section, we present the results of applying our methods to the two previously published sets of data introduced in the Methods section, the ESR set and the human orthologs of the Alarm Pheromone set. For both above data sets, we show the results from analyzing the genes using the biological process (BP) category of the gene ontology. These results will show 1) areas of agreement and difference between Benjamini-Hochberg and random resampling in evaluation of FDR, 2) how the assessment of significance of enrichment varies according to the particular database that is being probed, and 3) how the assessment of significance of enrichment varies according to the weight assigned to precision vs. recall.

### ESR Set (Environmental Stress Response, yeast)

#### Benjamini-Hochberg (BH)

Figure 2 shows the results of F-measure optimization on the ESR data based on FDR calculated by Benjamini-Hochberg (BH) method. As expected by their definitions, precision (F_0_) decreases with increasing *p*-value while recall increases with increasing *p*-value. F_0.5_ (precision-emphasized), F_1_ (precision and recall equally weighted) and F_2_ (recall-emphasized) all show relative maxima, providing a rational basis for assigning a threshold for significance. The horizontal scale is extended far enough to visualize the determination of the number of real positives. In the case of the up-regulated gene set, maximum F_1_ occurs at an uncorrected *p*-value close to 0.05. In the case of the down-regulated gene set however, it appears that a much more stringent cutoff would be appropriate.

**Figure 2.**
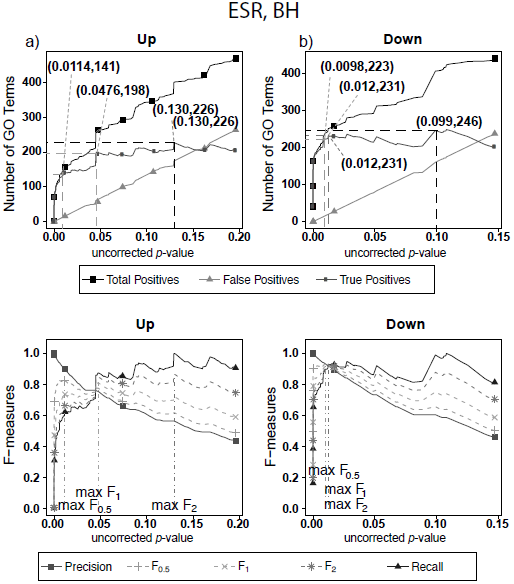
Number of positives and F-measure values for ESR set, BH-estimated FDR. a) Shows the number of enriched biological process Gene Ontology categories as a function of uncorrected p-value, the Benjamini-Hochberg number of false discoveries, and the projected true positives, namely the difference between the predicted positives and the false positives, for the upregulated ESR gene set. This panel is from the same data set at Figure 1. The number pairs in parenthesis are respectively (uncorrected *p*-value maximizing F_0.5_, number of true positives at that *p*-value), (uncorrected *p*-value maximizing F_1_, number of true positives at that *p*-value), (uncorrected *p*-value maximizing F_2_, number of true positives at that *p*-value), (uncorrected *p*-value maximizing true positives, number of true positives at that *p*-value) b) is the same as a) for the downregulated gene set. c) shows the F-measures computed from a) and d) the F-measures computed from b). Number of real positives, necessary to calculate recall (and therefore (F-measure)), is approximated by (predicted positives – false positives) _max_. The *p*-value at which the computed true positives are a maximum is 0.13 for up-regulated gene list (a) and at 0.099 for downregulated gene list. (b) The pairs of numbers in parenthesis in a) and b) indicate the *p*-value and number of returned GO terms at significant markers, specifically at maximum F_0.5_ (emphasizing precision), F_1_ (balanced emphasis between precision and recall), F_2_ (emphasizing recall), and Recall where we obtain an estimation of relevant elements by maximizing true positive).

#### Resampling

Figure 3 shows the results of F-measure optimization on the ESR data using resampling to calculate FDR. The false positives are calculated by average number of GO categories enriched in random sets. For the up-regulated set, all the F-measures optimize at much lower uncorrected *p*-values than do the F-measures calculated by the BH method. For the down-regulated set, resampling-calculated F_0.5_ is optimized at a lower uncorrected *p*-value than BH method while F_1_ and F_2_ are optimized at slightly higher uncorrected *p*-value.

**Figure 3.**
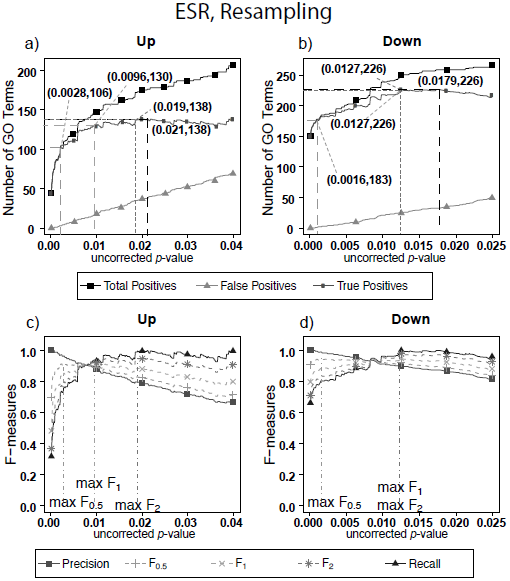
Number of positives and F-measure values for ESR set, Resampling-estimated FDR. A) Shows the number of enriched biological process Gene Ontology categories as a function of uncorrected p-value, the average number of enriched Gene ontology categories from the random set as the false positives, and the projected true positives, namely the difference between the predicted positives and the false positives, for the up-regulated ESR gene set. The number pairs in parenthesis are respectively (uncorrected *p*-value maximizing F_0.5_, number of true positives at that *p*-value), (uncorrected *p*-value maximizing F_1_, number of true positives at that *p*-value), (uncorrected *p*-value maximizing F_2_, number of true positives at that *p*-value), (uncorrected *p*-value maximizing true positives, number of true positives at that *p*-value) b) is the same as a) for the down-regulated gene set. c) shows the F-measures computed from a) and the F-measures computed from b). Number of real positives, necessary to calculate recall (and therefore (F-measure)), is approximated by (predicted positives – false positives) _max_. The *p*-value at which the computed true positives are a maximum is 0.021 for upregulated gene list (a) and 0.0179 for downregulated gene list. (b) The pairs of numbers in parenthesis in a) and b) indicate the p-value and number of returned GO terms at significant markers, specifically at maximum F_0.5_ (emphasizing precision), F_1_ (balanced emphasis between precision and recall), F_2_ (emphasizing recall), and Recall (where we obtain an estimation of relevant elements by maximizing true positive).

Comparing the results in Figure 2 and Figure 3 show that the optimum cutoff (as measured by maximum F_1_) varies widely, depending on the gene set to be tested and the method for assessing FDR. Using BH the optimum cutoff is .0476 for upregulated ESR and .012 for downregulated ESR. Using resampling, the optimum cutoff is.0096 for upregulated ESR and.0126 for downregulated ESR. Also, as expected, the optimum cutoff is relaxed when recall is emphasized (F_2_ instead of F_1_) and made more stringent when precision is emphasized (F_0.5_ instead of F_1_).

### Alarm Pheromone Set (human orthologs)

#### Benjamini-Hochberg (BH)

Figure 4 shows exactly the corresponding results as Figure 2, this time on the human orthologs to the honey bee alarm pheromone set. F-measures are maximized at much higher thresholds than for the ESR set. The difference in optimal F-measure is largely due to the different shapes of the recall curves. For the ESR set, precision drops significantly more rapidly with increasing uncorrected *p*-value than does the AP set. Therefore, a higher uncorrected *p*-value can be used for the latter set with essentially the same degree of confidence.

**Figure 4.**
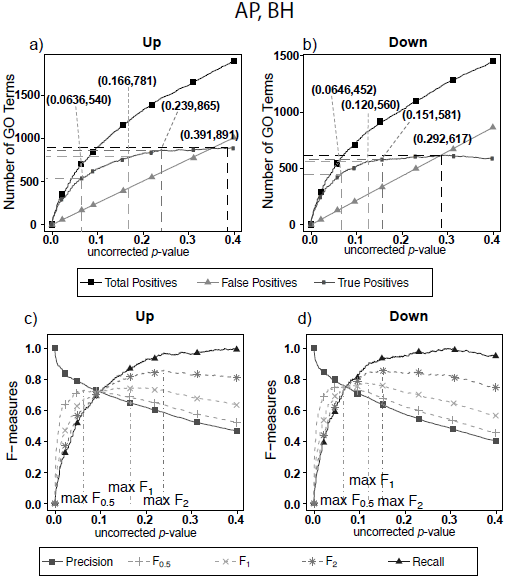
Number of positives and F-measure values for Alarm Pheromone set, BH-estimated FDR. a) shows the number of enriched biological process Gene Ontology categories as a function of uncorrected *p*-value, the Benjamini-Hochberg number of false discoveries, and the projected true positives, namely the difference between the predicted positives and the false positives, for the upregulated alarm pheromone human orthologs gene set. The number pairs in parenthesis are respectively (un-corrected *p*-value maximizing F_0.5_, number of true positives at that *p*-value), (uncorrected *p*-value maximizing F_1_, number of true positives at that *p*-value), (uncorrected *p*-value maximizing F_2_, number of true positives at that *p*-value), (uncorrected *p*-value maximizing true positives, number of true positives at that *p*-value) b) is the same as a) for the down-regulated gene set. c) shows the F-measures computed from a) and d) the F-measures computed from b). Number of real positives, necessary to calculate recall (and therefore (F-measure)), is approximated by (predicted positives – false positives) _max_. The *p*-value at which the computed true positives are a maximum is 0.391 for upregulated gene list (a) and at 0.292 for downregulated gene list. (b) The pairs of numbers in parenthesis in a) and b) indicate the p-value and number of returned GO terms at significant markers, specifically at maximum F_0.5_ (emphasizing precision), F_1_ (balanced emphasis between precision and recall), F_2_ (emphasizing recall) and Recall (where we obtain an estimation of relevant elements by maximizing true positive).

#### Resampling

Figure 5 shows the number of GO categories and F-measures for the alarm pheromone set human orthologs using resampling method. The resampling method have found more false positives than BH, and therefore the precision is much lower than the precision calculated from BH, and the F-measures are optimized at lower uncorrected *p*-values than the F-measures calculated from BH.

**Figure 5.**
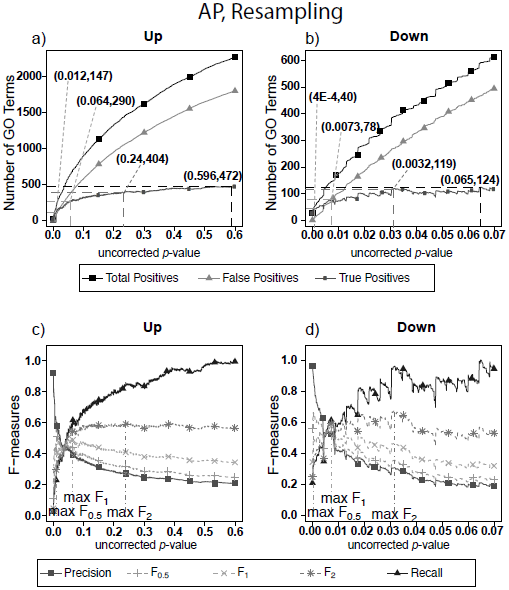
Number of Positives and F-measure values for AP set, Resampling-estimated FDR. The figure shows the number of enriched biological process Gene Ontology categories as a function of uncorrected *p*-value, the average number of enriched Gene ontology categories from the random set as the false positives, and the projected true positives, namely the difference between the predicted positives and the false positives, for the up-regulated alarm pheromone human orthologs gene set. b) is the same as a) for the down-regulated gene set. c) shows the F-measures computed from a) and d) the F-measures computed from b). Number of real positives, necessary to calculate recall (and therefore (F-measure)), is approximated by (predicted positives – false positives) _max_. The *p*-value at which the computed true positives are a maximum is 0.596 for upregulated gene list (a) and at 0.065 for downregulated gene list. (b) The pairs of numbers in parenthesis in a) and b) indicate the *p*-value and number of returned GO terms at significant markers, specifically at maximum F_0,5_ (emphasizing precision), F_1_ (balanced emphasis between precision and recall), F_2_ (emphasizing recall), and Recall (where we obtain an estimation of relevant elements by maximizing true positive).

From the above Figures 2-5, we can note the stepped structure in the number of enriched GO categories. The stepped structure lies in the fact that the number of genes associated with any GO category, in the test set or reference set, must be an integer with limited number of choices. Therefore, the uncorrected *p*-values calculated would be in a discrete set instead of a continuum. Consequently, the number of positives as a function of *p*-values increases in a stepped way. As a result, the F-measures derived from the number of GO categories have spikes. But as our graphs have demonstrated, the optimal F-measures reflect the different weights on precision and recall despite the spikes.

Comparing the results in Figures 4 and 5 shows that, for the AP gene sets as for the ESR gene sets, the optimum cutoff threshold is different for the upregulated and downregulated gene sets and also is different when BH is used to determine the FDR as compared to resampling.

### Comparison of F-Measure with MCC for Optimization of Threshold Choice

As indicated in the section on methods, a widely used alternative to the F-measure for optimization is the Matthews Correlation Coefficient (MCC) which, unlike the F-measure, gives equal weight to negative as well as positive identifications. Figure 6 shows MCC optimization for exactly the same data set (ESR) and False Discovery Rate determination (Resampling) as in Figure 5. The most important lesson from this Figure is that the uncorrected p-value that maximizes MCC_1_ is the same as the uncorrected p-value that maximizes F_1_. Inspection of the formulas reveals the reason. The divergence between MCC and F-measure occurs only when the false negatives are a significant fraction of the total negatives.

**Figure 6.**
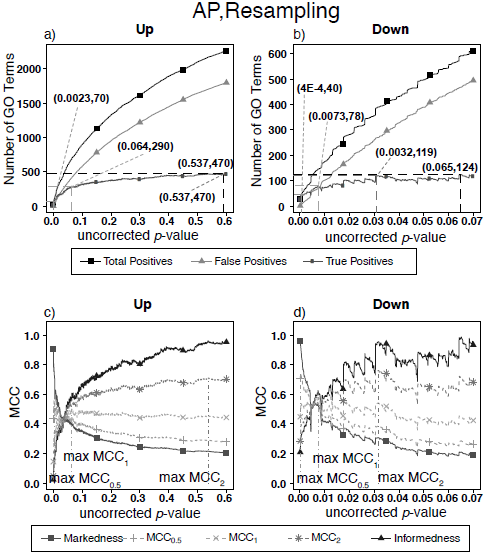
Number of Positives and MCC-measure values for AP set, Resampling-estimated FDR. This figure is the same as Figure 5 except that the optimization to determine significance-insignificance threshold is Matthews Correlation Coefficient (MCC) rather than F-measure. Note that the uncorrected p-value threshold for optimum MCC_1_ is the same as for F_1_. Examination of the expressions for the two quantities shows that the reason for the convergence is that in this case the number of false negatives is very small compared to the number of total and true negatives, so the fractional variation in true negatives is very small. This is true for all the data sets.

Since there are tens of thousands of terms in the gene ontology database this condition does not pertain to our situation, so optimization of the F-measure is an adequate strategy. However, we agree with Powers ^26^ that optimization of the MCC is the more universally correct strategy.

### Comparison of FDR (False Positive) Calculation by Benjamini-Hochberg (BH) and Resampling

In the previous section, we have demonstrated how to use F-measure optimization to obtain a flexible threshold based on whether precision or recall is more heavily weighted by the researcher. In that section the FDR is calculated but not shown explicitly. The present section explicitly compares the FDR as calculated by the BH method and by random resampling. In each case the random resampling FDR is computed based on the average of 50 randomly sampled null sets of the same size as the test set. Figure 7 shows that for the ESR set, the BH method and resampling estimate similar FDR at low *p*-value. As the threshold increases, the BH method estimates lower false discovery rate, and therefore higher precision, than the resampling method at the same uncorrected *p*-value. By contrast, for the Alarm Pheromone set, the BH method estimates lower FDR than resampling.

**Figure 7.**
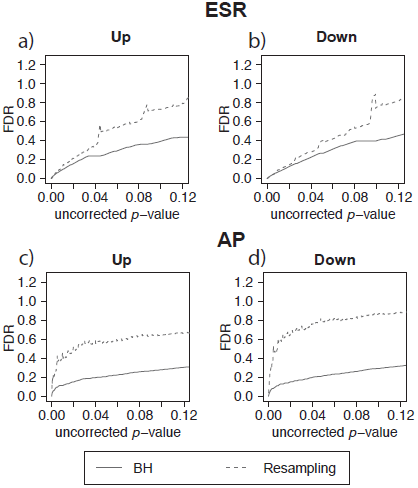
False discovery rate comparison. False discovery rate estimated by Benjamini-Hochberg (solid curve) and Resampling (dashed curve) for the ESR set and Alarm Pheromone set. Figure 7 compares the number of false discovery rate calculated by Benjamini-Hochberg (solid) and Resampling (dashed) in each set: a) up-regulated ESR, b) down-regulated ESR, c) up-regulated Alarm Pheromone set, and d) down-regulated Alarm Pheromone set. Generally, resampling has found higher false discovery rate than Benjamini-Hochberg. At low *p*-values, the BH and resampling methods get similar estimation of false discovery rate for the ESR set.

To further evaluate the methods, we carried out multiple runs using random (null) sets as test sets. In this case, the FDR should in principle be 1, for any uncorrected *p*-value. The results of this test are shown in Figure 8a, where for each segment of *p*-values (bin size = 0.0001) we show the mean plus/minus the standard deviation. The resampling method passes the test on the average, but the results are noisy. The BH method systematically underestimates FDR. Figure 7b shows that the noise in the resampling method results in Figure 7a are largely due to the variation in the random null sets, and that the noise level in using random resampling for real data is acceptably low.

**Figure 8.**
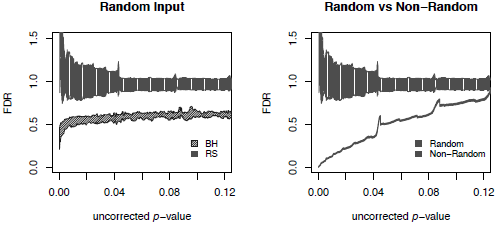
Comparison of different FDR calculation method on accuracy and convergence. a) Comparison of BH and Resampling on random “test” sets. At each *p*-value (*p*-values binned at intervals of.0001), the mean and standard deviation are calculated and plotted as shown. The random test sets consist of 281 yeast genes, against the background of the entire yeast genome. For each of the methods 50 test sets were used and the mean plus/minus standard deviation plotted as shown. Resampling hits the mark on the average but with substantial noise, while BH systematically underestimates FDR. b) Evaluation of resampling convergence on a real data set, ESR upregulated considered in this paper. This set is run against five different ensembles of null sets, each ensemble containing 100 null sets. The mean and standard deviation are plotted and compared to the results from the random test sets. It is seen that the noise of the resampling method on a real data set is acceptable.

## Statistical Summary of Results from Different Threshold Criteria

Table 1 shows the statistical summary of using all different criteria for the distinction between significant and non-significant enrichment. Notable features of this table include: 1) Variation of the threshold within the range explored in this study made relatively little statistical difference for the ESR set. Over the entire range of thresholds, both the precision and the recall for the ESR set are good, and the number of terms returned does not change very much. 2) Variation of the threshold within the range explored in this study makes a very large difference in the results of the AP set. For the most stringent choice of threshold, the precision is high, but the recall is quite low. Relaxing the threshold improves the recall, but at a cost to the precision, so there is a distinct tradeoff between precision and recall, and 3) We discovered that optimizing F_1_ is exactly equivalent to optimizing the Matthews correlation coefficient. F_.5_ is optimized at a lower uncorrected p-value than F_1_ while F_2_ is optimized at a higher p-value, and the same pattern is seen for MCC.

**Table 1.**
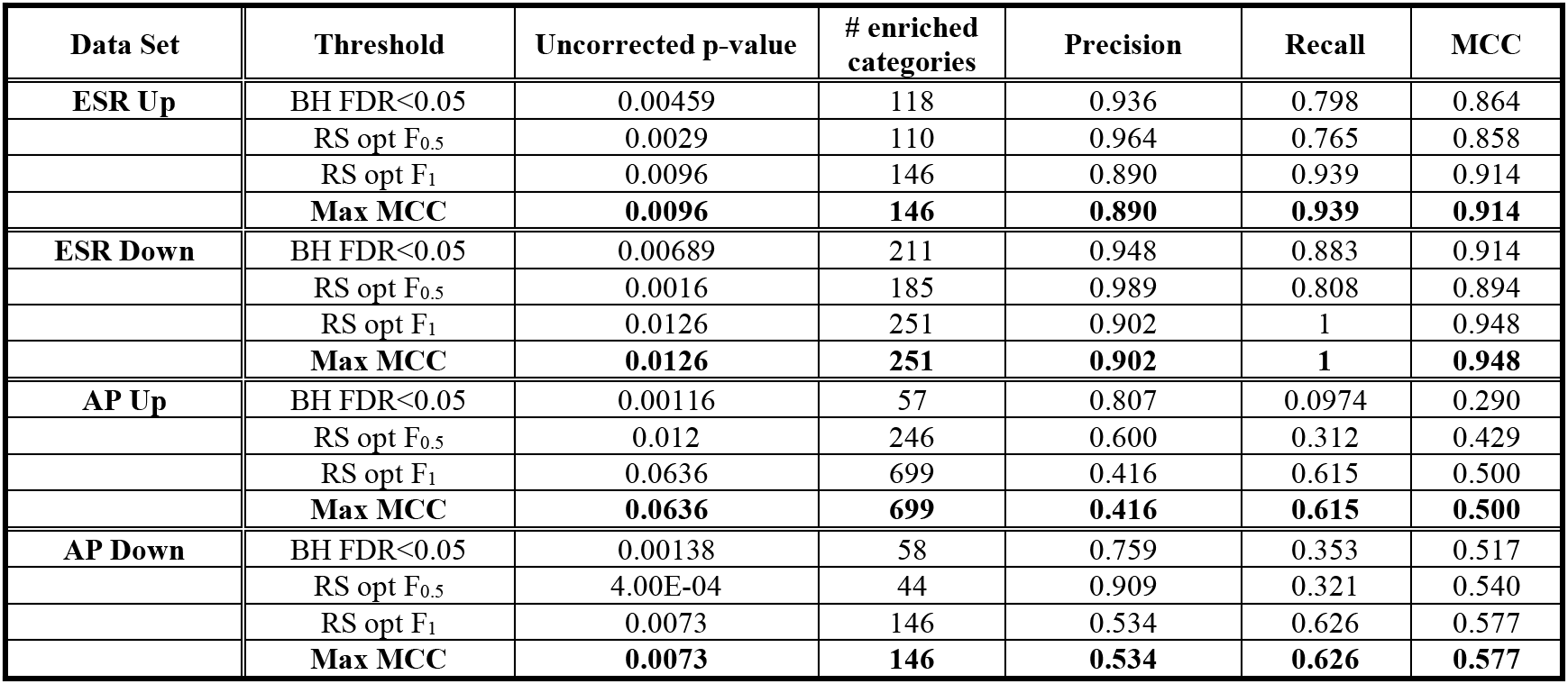
Precision, Recall, and Matthews Correlation Coefficients (MCC) at thresholds BH FDR<0.05, Resampling optimal F_0.5_, and Resampling optimal F_1_. For the four data sets examined, we have found that optimal F_1_ is the position that MCC reaches maximum. This correspondence between optimum F_1_ and optimum MCC was unanticipated but emerged from independent calculation of both quantities. For the ESR set, the MCC is high for all thresholds. For AP set, MCC is relatively low, and the MCC for BH FDR<0.05 is the lowest.

### Identity of Enriched Terms Using Different Threshold Criteria

#### A. Higher order relatively general terms

The enriched GO terms are categorized by their parent terms, 1^st^ order parent being direct children of the root term “Biological Process” (GO:0008150), 2^nd^ order parent being direct children of the 1^st^ order parent terms. Each enriched GO term is traced back to the root by the shortest route. Tables 2 through 5 below provide an outline of the complete gene ontology results by showing the high order terms that are either themselves enriched according to the described criteria or have child terms enriched, or both. In each case the results from three different thresholds are shown, BH FDR<.05, optimum F_.5_, and optimum F_1_. The most striking pattern is that for the ESR sets (Tables 2 and 3), modifying the threshold within the parameters of this paper did not change the identity of the putatively enriched higher order terms very much. However, for the AP sets (Tables 4 and 5), relaxing the threshold caused a substantial increase in the number of high order terms judged to be putatively significant. However, from Table 1 is it seen that the precision (confidence) of the additional terms for the AP sets is substantially lower than for the terms returned using the most stringent threshold. Thus, for the AP set we clearly see that we can’t simultaneously have high precision and high recall. We must trade one for the other.

**Table 2.** ESR, Up-regulated Set. Each row corresponds to a 1^st^ order Parent Terms of enriched GO categories of ESR set, Up regulated genes. The three numbers in parentheses reflect the total number of terms in the Parent family (Parent plus children). We found no difference in the high order terms between BH FDR<.05 and F_.5_ However the developmental process parent term (labeled with “**”) emerges when the threshold is increased to optimal resampling F_1_. The groupings as defined by the parent terms do not change very much, but the number of more specific child terms increases moderately.

| GO ID | Parent Term | Minimum raw $p$-value of child terms |
| --- | --- | --- |
| GO:0008152 | Metabolic Process (80,85,100) | 3.40E-13 |
| GO:0050896 | response to stimulus (22,23,26) | 7.40E-13 |
| GO:0065007 | biological regulation (4,5,7) | 9.00E-05 |
| GO:0009987 | cellular process (4,5,13) | 0.00035 |
| **\*\*GO:0032502** | developmental process (0,0,1) | 0.00589 |

**Table 3.** ESR, Down-regulated Set. 1^st^ order Parent Terms of enriched GO categories of ESR set, down regulated genes. For this data set the optimum was more stringent than the BH FDR <.05. The term “response to stimulus” (labeled with “*” does not meet the optimum F_.5_ criterion but does for the other two criteria. The numbers in the parentheses refer to the numbers of enriched terms in each parent category, ordered from low to high. As with the up-regulated genes, relaxing the threshold did not change the parent terms much, but did increase the number of more specific child terms moderately.

| GO ID | Parent Term | Minimum raw $p$-value of child terms |
| --- | --- | --- |
| GO:0008152 | Metabolic Process (120,139,168) | 1.00E-30 |
| GO:0009987 | Cellular process (6,6,7) | 1.00E-30 |
| GO:0071840 | Cellular component organization or biogenesis (31,32,36) | 1.00E-30 |
| GO:0051179 | Localization (21,22,22) | 5.20E-28 |
| GO:0065007 | biological regulation (7,11,15) | 3.20E-12 |
| **\*GO:0050896** | response to stimulus (0,1,2) | 0.00357 |

**Table 4.** 1^st^ order Parent Terms of enriched GO categories of AP set, Up regulated genes. The terms with “*” appears when the threshold is increased from BH FDR<0.05 (uncorrected p-value<0.00116) to optimal resampling F_0.05_-measure (uncorrected p-value<0.012). Terms with “**” emerges when the threshold is increased to that for optimal resampling F_1_(uncorrected p-value<0.0096). The number in the brackets refers to the number of enriched terms within each parent category at each threshold, ordered from low to high. Unlike the ESR sets, for this data set relaxing the threshold caused significantly greater returns in both general terms and their children.

| GO ID | Parent Term | Minimal raw $p$-value of child terms |
| --- | --- | --- |
| GO:0009987 | Cellular process (13,36,96) | 1.10E-10 |
| GO:0050896 | Response to stimulus (57,71,119) | 1.40E-08 |
| GO:0065007 | Biological regulation (28,113,288) | 4.30E-05 |
| GO:0008152 | Metabolic process (9,44,113) | 5.00E-05 |
| GO:0032502 | Developmental process (1,9,33) | 0.00043 |
| GO:0071840 | cellular component organization or biogenesis (1,6,12) | 0.00102 |
| *GO:0051179 | Localization (0,8,37) | 0.00138 |
| *GO:0022414 | reproductive process (0,2,7) | 0.00192 |
| *GO:0002376 | immune system process (0,2,8) | 0.00504 |
| *GO:0032501 | multicellular organismal process (0,5,19) | 0.00509 |
| *GO:0040011 | Locomotion (0,1,2) | 0.00932 |
| **GO:0051704 | multi-organism process (0,0,11) | 0.02 |
| **GO:0008283 | cell proliferation (0,0,2) | 0.02962 |

**Table 5.** 1^st^ order Parent Terms of enriched GO categories of AP set, Down regulated genes. The terms with “*” disappears when the threshold is decreased from BH FDR<0.05 (uncorrected p-value<0.00138) to optimal resampling F_0.05_-measure (uncorrected p-value<4.00E-4). Terms with “**” emerges when the threshold is increased at optimal resampling F_1_(uncorrected p-value<0.0073). The number in the brackets refers to the number of enriched terms at each threshold, low to high. Unlike the ESR sets, for this set relaxing the threshold caused substantial increases in the putative enriched categories at both the general level and the more specific child level.

| GO ID | Description | Minimal $p$-value of child terms |
| --- | --- | --- |
| GO:0008152 | Metabolic Process (40,7,25) | 3.20E-08 |
| GO:0009987 | cellular process (3,4,13) | 7.00E-06 |
| GO:0071840 | cellular component organization or biogenesis (1,0,5) | 7.90E-06 |
| *GO:0051179 | Localization (0,3,16) | 0.00052 |
| **GO:0065007 | biological regulation (0,0,15) | 0.00145 |
| **GO:0050896 | response to stimulus (0,0,7) | 0.00174 |
| **GO:0022414 | reproductive process (0,0,1) | 0.00441 |
| **GO:0051704 | multi-organism process (0,0,1) | 0.00441 |
| **GO:0032501 | multicellular organismal process (0,0,3) | 0.00441 |
| **GO:0032502 | developmental process (0,0,1) | 0.00534 |

#### B. Relatively Specific Terms

Specific, or “child” terms returned in these calculations are too numerous to delineate completely in the body of the paper. They are instead provided in the spreadsheet “AllGOTermsInTree_Final(supplementary material 1)” Separate tabs delineate the returns from ESR upregulated, ESR downregulated, AP upregulated, and AP downregulated. Each entry in the spread sheet is color coded with the code given in the tab labeled “color coding”. Entries that are shaded are either primary or secondary (more general) classes, which will also be shown in Table 1. Entries colored in black appear at “standard” threshold: BH FDR<0.05. Entries colored in blue emerge at the threshold determined by optimal F_0.5_. For AP Up, the standard threshold is the most stringent while for all other sets, the optimal F_0.5_ is the most stringent. Entries colored in red first emerge at the least-stringent threshold for that data set, which corresponding to optimal F_1_. The format of the spreadsheet for each of the data sets is as follows: Column A is the identifying number of the GO class that is returned as significant, column B is the name of that class, and column C is the raw enrichment *p*-value for that class. Column D is non-zero only for the rows belonging to primary or secondary GO classes (which are shown explicitly in Tables 2-5 for the four data sets). The numerical value in column D represent the smallest uncorrected *p*-value of all the classes under the primary or secondary class shown in that row. The spread sheet is organized to be sectioned off according to primary or secondary classes. To illustrate the sectioning, under the “AP up” is the primary class “cellular process” and immediately under that the secondary class “protein folding”. This is followed by more specific classes under “protein folding” such as “chaperone-mediated protein folding” and others. The columns E and farther to the right are GO numbers representing the lineage of the particular term in that row starting with the primary class and continuing to the particular term in that row.

Because the trade-offs with varying threshold are most clear with the AP sets, we select those now for discussion. One biologically interesting feature emerging from varying the threshold consists of the more specific GO classes emerging from general classes already identified with a more stringent threshold. For example, in the “AP up” set “protein folding” was identified as a secondary class of interest by virtue of a very strong enrichment score. On relaxing the threshold more specific “child” classes emerged, such as “chaperone cofactor-dependent protein folding”, “endoplasmic protein folding”, and others. While these more specific classes are identified with less confidence than the overall “protein folding” class they are subsumed into, they do provide the most likely subclasses within protein folding to be biologically meaningful. Similarly, under the secondary class of “signal transduction” more specific subclasses such as “ER-nucleus signaling pathway”, “stress-activated MAPK cascade” and others emerge with modest threshold relaxation. This pattern is seen throughout the spreadsheet. Relaxing the threshold provides not only improved recall, but improved specificity, which will help in biological interpretation of GO enrichment results.

#### C. Summary

In general, when thresholds are varied, a tradeoff can plainly be seen between precision and recall. When looking at the specific GO classes that are returned at different choices of threshold a second tradeoff emerges, between generality and specificity. As threshold is relaxed some more general terms are revealed, but the greater effect is that more specific terms are revealed within general terms that were suggested at more stringent thresholds. These specific terms can help to provide a more focused interpretation of the biological results.

## Conclusions

In this work, we have addressed two issues with the commonly used methods in the GO enrichment analysis: the relationship between resampling vs. Benjamini-Hochberg theory for estimating false discovery rate, and the arbitrariness of the threshold for significance.

To consider resampling vs. Benjamini-Hochberg we made five independent comparisons. Four consisted of upregulated and downregulated genes separately for two different animal experiments. The fifth was an array of random gene lists (null sets). For the yeast ESR sets the two methods gave almost the same results for uncorrected p-value<.04 but diverged substantially for more relaxed *p*-values, with the BH underestimating the FDR. For the honeybee AP set the BH method underestimated the FDR significantly at all uncorrected *p*-values. For the random or null sets, we know that the correct FDR is 1, because there is no significance to the results. Yet for the null sets the BH method produced FDR<1 by a large margin for the full range of uncorrected *p*-values. By contrast the resampling method, although noisy, does not systematically deviate from 1 in its prediction of FDR for the null sets.

It is of interest to consider why the BH method, while very useful and successful in some cases, sometimes fails. It is understood that the method will always work when the true inferences are independent. Strictly speaking, this will not be true of Gene Ontology data since many genes belong in multiple Gene Ontology categories. However, Benjamini and Yekutieli^32^ showed that the method was still valid for dependent hypotheses provided that the related hypotheses that failed the null test showed positive regression of likelihoods. Consideration of the tree-like structure of Gene Ontology data^33^ shows that this is true to a great extent. The branches of the tree-like structure clearly show positive regression within each branch; if a child category is enriched a parent is more likely to be enriched, and vice versa. Thus, as long as the enriched classes fall along a few well-delineated branches of the Gene Ontology tree structure, BH will work well. This appears to be largely the case for the yeast ESR set at relatively stringent *p*-values, in which the experimental intervention activated well-defined and annotated pathways. Thus, for relatively stringent cutoffs the BH FDR works well for this data set. However, some genes are members of categories in multiple branches, compromising the positive regression criterion. In the ESR set at relatively relaxed thresholds, and for the AP set at all thresholds, many Gene Ontology categories in different branches but with overlapping gene membership are represented in the returned categories, so that both independence and the positive regression criterion are violated. These considerations tell us why BH fails dramatically for the completely null sets. Neither independence nor positive regression are satisfied, except sometimes completely accidentally. For the issue of the arbitrariness of the threshold, we introduced optimization of F-measures so that both type I and II errors are considered. Unlike arbitrarily applied threshold of BH FDR<0.05 or uncorrected *p*-value<0.01 for any data set, the F-measure optimization approach provides a flexible threshold appropriate to the nature of the data set and the research question. If the data set is high in noise-to-signal ratio and the penalty for letting in false positive is high, we can choose to optimize F-measures weighing more on precision. If the data set fails to show much enrichment by commonly-applied methods, we can relax the threshold and extract the best information indicated by F-measure optimization.

A concern is that, because of the nature of the problem, we were forced to use a heuristic (albeit reasonable) method to estimate the false negatives, essential for calculating recall. We judge that this concern is more than offset by the advantage of enabling the replacement of an arbitrary threshold with F-measure optimization.

We found that for the particular class of problems dealt with in this paper the F-measure is as appropriate an optimization criterion as the Matthews Correlation Coefficient.

By examination of the specific GO categories that are returned by our analysis, we find that relaxing the threshold, we see revealed the most likely specific subcategories within the general categories that are revealed at the most stringent threshold. Thus, varying the threshold not only reflects the tradeoff between precision and recall, but also between generality and specificity.

In the supplementary material we present the spreadsheet **“AllGOTermsInTree_Final”,** which shows all the specific GO terms returned in the work described in this paper. Also, in the supplementary material, we present our automatic pipeline integrating TopGO with resampling and analyzing functions to carry out the whole process of resampling, enrichment analysis, F-measure calculation, and representing results in tables and figures. The pipeline also includes a GOstats ^15^ module for easy analysis of under-represented terms and a STRINGdb ^34^ module for KEGG pathway terms. As demonstrated, the pipeline can also calculate analytical FDR including, but not limited to, the BH method. In summary, we suggest replacing a fixed *p-*value for assigning a threshold in enrichment calculations with an optimal F-measure, which incorporates the well-established and well-defined concepts of precision and recall.

## Abbreviations

GO: gene ontology
FDR: false discovery rate
BH: Benjamini-Hochberg method
BP: biological process
CC: cellular component
MF: molecular function
BY: Benjamini-Yekutieli method
ESR: environmental stress response genes
AP: honey bee genes in response to Alarm Pheromone, human orthologs.

## Ethics approval and consent to participate

N/A

## Consent for publication

N/A

## Availability of data and material

Computer codes available in Supplementary Material

## Authors contributions

WG and ZF both did parts of the calculation and worked together to initially develop the automated pipeline. WG did final enhancement and debugging. EJ suggested the overall direction of the work. WG wrote the first draft of the manuscript. All three authors worked on refining the manuscript

## Acknowledgements

We gratefully acknowledge useful discussions with especially Dr. Enes Kotil and Santiago Nunez-Corrales and also other members of our research group in the Beckman Institute. Professor Saurabh Sinha of our Computer Science Department provided useful discussions and wise advice.

## Additional Files

Additional file 1--- AllGOTermsInTree_Final.xlsx

This is the spreadsheet showing all enriched terms at thresholds: BH FDR<0.05, optimal F_0.5_, and optimal F_1_. The terms are arranged by the primary and second-order parent terms.

Additional file 2 --- pipelinemanual.docx

“A TopGO- and GOstats-based automated pipeline for GO enrichment analysis using F-measure optimization based on resampling and traditional calculation”

This is a word document giving detailed description of how to run the pipeline for resampling or analytical FDR calculation and obtain thresholds maximizing F-measures

Additional file 3 --- pipeline.gz

This file contains source codes of the pipeline and the ESR and AP data sets for demo runs.

